# Selective transport of fluorescent proteins into the phage nucleus

**DOI:** 10.1101/2020.11.25.398438

**Authors:** Katrina T. Nguyen, Joseph Sugie, Kanika Khanna, MacKennon E. Egan, Erica A. Birkholz, Jina Lee, Christopher Beierschmitt, Elizabeth Villa, Joe Pogliano

**Affiliations:** Division of Biological Sciences, University of California San Diego, La Jolla, CA 92093, USA

## Abstract

Upon infection of *Pseudomonas* cells, jumbo phages 201Φ2-1, ΦPA3, and ΦKZ assemble a phage nucleus. Viral DNA is enclosed within the phage-encoded proteinaceous shell along with proteins associated with DNA replication, recombination and transcription. Ribosomes and proteins involved in metabolic processes are excluded from the nucleus. RNA synthesis occurs inside the phage nucleus and messenger RNA is presumably transported into the cytoplasm to be translated. Newly synthesized proteins either remain in the cytoplasm or specifically translocate into the nucleus. The molecular mechanisms governing selective protein sorting and nuclear import in these phage infection systems are currently unclear. To gain insight into this process, we studied the localization of five reporter fluorescent proteins (GFP^+^, sfGFP, GFPmut1, mCherry, CFP). During infection with ΦPA3 or 201Φ2-1, all five fluorescent proteins were excluded from the nucleus as expected; however, we have discovered an anomaly with the ΦKZ nuclear transport system. The fluorescent protein GFPmut1, expressed by itself, was transported into the ΦKZ phage nucleus. We identified the amino acid residues on the surface of GFPmut1 required for nuclear targeting. Fusing GFPmut1 to any protein, including proteins that normally reside in the cytoplasm, resulted in transport of the fusion into the nucleus. Although the mechanism of transport is still unknown, we demonstrate that GFPmut1 is a useful tool that can be used for fluorescent labelling and targeting of proteins into the ΦKZ phage nucleus.

## Introduction

Protein targeting within a cell is essential in all organisms. Generally, eukaryotes use a sorting sequence to target proteins to specific organelles, such as a nuclear localization signal to send proteins to the nucleus or an N-terminal signal peptide to target proteins to the endoplasmic reticulum. These signal sequences are usually highly conserved, even among different species (1, 2). Though bacterial cells lack the membrane-bound organelles of eukaryotes, they still utilize a number of protein sorting strategies to target proteins either extracellularly or to specific intracellular locations (3–5). For example, secretion of unfolded proteins from the cytoplasm requires a signal sequence, which directs proteins to the SecYEG pore where secretion is powered by the ATPase SecA and the proton motive force (4, 6). In contrast, the TatA system exports fully folded proteins across the cytoplasmic membrane after recognizing a pair of arginine residues at the C-terminus (5). The Sec and Tat pathways are highly conserved in all domains of life (3). In addition to these general secretory systems, many additional systems (Type I - VI) transport specific cargo across the inner and outer bacterial membranes (3). These transport systems all utilize a beta-barrel channel that spans the membrane but are widely divergent in most other aspects (3).

Protein targeting is essential for establishing and maintaining subcellular organization as well as for viral replication. We recently described the phage nucleus assembled by jumbo phages 201Φ2-1 (7, 8), ΦPA3 (9), and ΦKZ (10) in *Pseudomonas* cells (11, 12). In the early stages of infection, the phage assembles a nucleus-like structure in the cell and positions it at midcell using a dynamic bipolar tubulin-based spindle (11–16). Phage proteins synthesized by bacterial ribosomes in the cytoplasm appear to be sorted to specific subcellular destinations based on their biological functions. Much like in a eukaryotic cell, proteins involved in DNA replication, repair, and transcription localize inside the nucleus, while proteins involved in metabolic processes and protein synthesis localize in the cytoplasm outside the nucleus (11, 12). Time-lapse microscopy experiments show that phage proteins, expressed in our heterologous system, are synthesized before phage are introduced, then accumulate in the nucleus as infection occurs, suggesting that a mechanism exists for posttranslational nuclear protein transport (12). However, no known eukaryotic nuclear localization signals or bacterial sorting sequences were encoded by the phages. In addition, we have not identified any homology to bacterial transporters or nuclear pore proteins in the phage genomes. The mechanisms of protein sorting and intracellular transport are still unknown.

One of the barriers to understanding the details of *Pseudomonas* jumbo phage replication is the inability to specifically target proteins, such as gene editing enzymes or other effectors, to the phage nucleus versus the cytoplasm. Here, we report a technique for targeting proteins into the ΦKZ nucleus. Although the nucleus of ΦKZ appears to be largely similar to that of phages ΦPA3 and 201Φ2-1, surprisingly, we found that it imports the fluorescent protein GFPmut1, but not any of the other tested fluorescent proteins. In addition, any protein fused to GFPmut1 also localized to the ΦKZ nucleus. Thus, we have serendipitously discovered a reliable method for delivering specific proteins into the ΦKZ nucleus.

## Results

During comparative protein localization of cells infected with one of three different phages (ΦPA3, ΦKZ, 201Φ2-1), we noticed a discrepancy in localization of the fluorescent proteins themselves. All fluorescent protein controls (GFPmut1, GFP^+^, sfGFP, mCherry, and CFP) were localized in the cytoplasm of ΦPA3 and 201Φ2-1 as expected (Fig 1A). Four of these proteins, GFP^+^, sfGFP, mCherry, and CFP also localized in the cytoplasm of ΦKZ infected cells (Fig 1A). However, GFPmut1 localized inside the ΦKZ nucleus even though it was excluded by the nucleus of 201Φ2-1 in *P. chlororaphis* and that of ΦPA3 in *P. aeruginosa* (Fig 1B). Our results suggest that differences exist among the fluorescent proteins that affect their ability to be transported into the ΦKZ nucleus.

**Fig 1:**
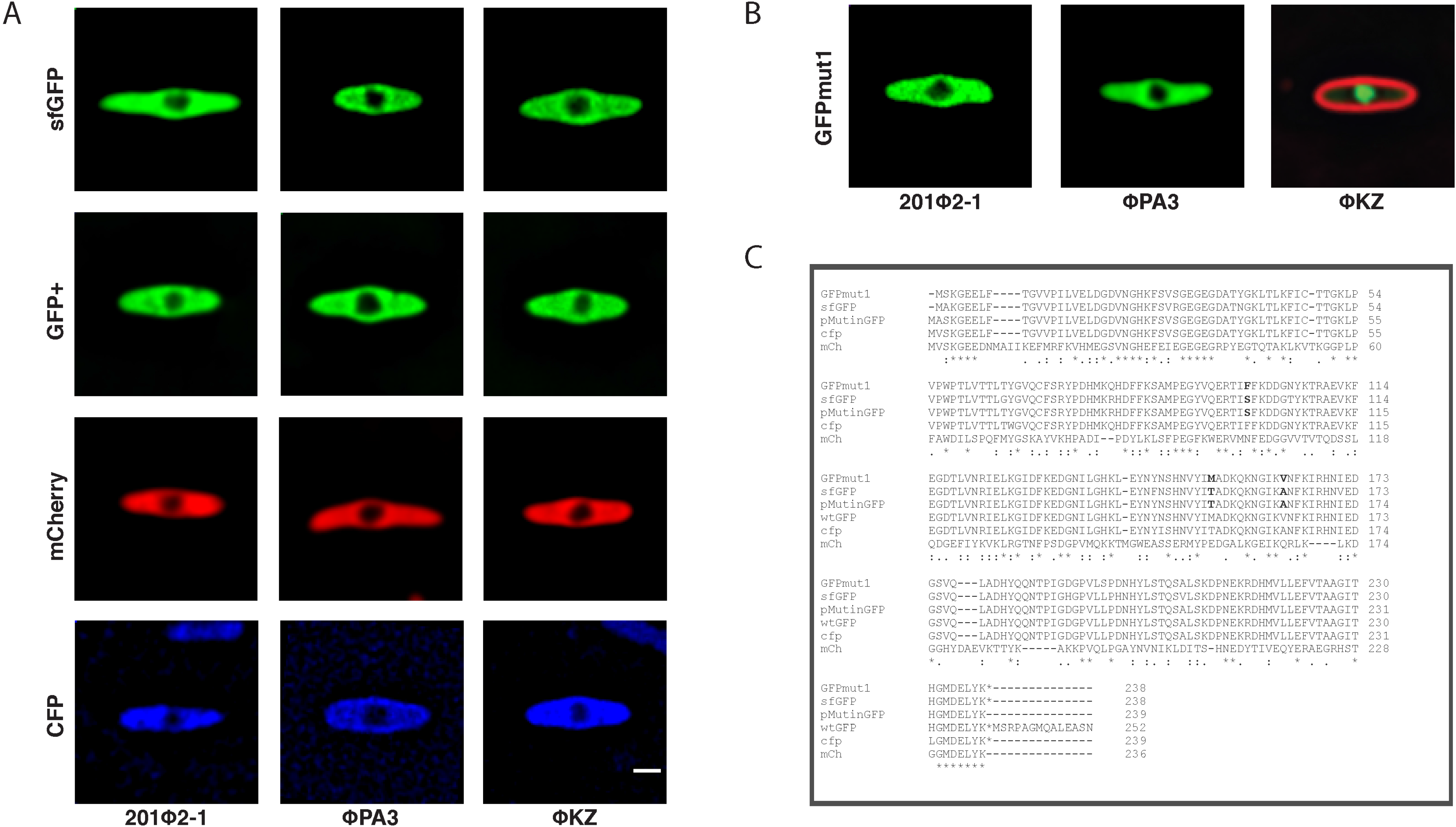
Fluorescent protein localization during phage infection. Most fluorescent proteins localize to the bacterial cytoplasm and are excluded by the phage nucleus but GFPmut1 is transported into the ΦKZ nucleus. Scale bar = 1 micron. A. SfGFP, GFP^+^, mCherry, and CFP are excluded by the phage nucleus in *P. chlororaphis* cells infected with 201Φ2-1 and *P. aeruginosa* cells infected with ΦPA3 or ΦKZ. B. GFPmut1 localizes inside the ΦKZ phage nucleus but is excluded from both the 201Φ2-1 nucleus and ΦPA3 nucleus. C. Alignment of fluorescent protein sequences showing key differences in bold letters. Key differences occur at F99, M153, and V163 of GFPmut1 compared to other fluorescent proteins.

We reasoned that studying nearly identical fluorescent proteins with strikingly different localizations might provide insights into nuclear targeting. Comparison of the protein sequences (17–21) of these fluorescent proteins revealed several amino acid differences that could be responsible for the discrepancy in localization of GFPmut1 (Fig 1C). We identified three amino acids where GFPmut1 differed from the other proteins tested: F99, M153, V163 (Fig 1C, bold letters, Fig S1) (17, 19, 22). In the 3-dimensional protein structure of GFP, phenylalanine (F99) and methionine (M153) are both surface exposed, extending outward from one face of the GFP, while valine (V163) is along the same surface but facing inward toward the beta barrel (Fig 2A) (19, 22–24). To determine which of these mutations might influence import into ΦKZ, we used site-directed mutagenesis to individually mutate each amino acid of GFPmut1 to that found in alternate versions of GFP, specifically those that remained in the cytoplasm during infection (sfGFP and GFP^+^) (17, 19, 25). At 60 minutes post-infection, GFPmut1 with the V163A mutation (n=82) retained the same phenotype as the unaltered GFPmut1 (n=111), localizing to the nucleus (Fig 2B, 2C). However, the F99S mutation completely changed the phenotype so that the fluorescent protein was localized in the cytoplasm and excluded by the nucleus in 100% of cells (n=177) (Fig 2B, 2C). The M153T mutation partially altered GFP localization creating a mixed phenotype among cells in the population (n=115) which we quantitated by plotting normalized pixel intensity profiles of fluorescence signals along a line through the center of the long axis of the cell (Fig 2C). Individual tracings (Fig 2C, blue) show there is significant variability among the population of M153T cells, ranging from fully excluded (minimum intensity at the center) to fully nuclear localized (maximum intensity at the center). In contrast, both the individual tracings (Fig 2C) and average tracings (Fig 2C, D) of GFPmut1 and V163A were fully imported while F99S was fully excluded. These results suggest that the amino acids on the surface of GFPmut1 contribute to its selective import into the ΦKZ nucleus.

**Fig 2:**
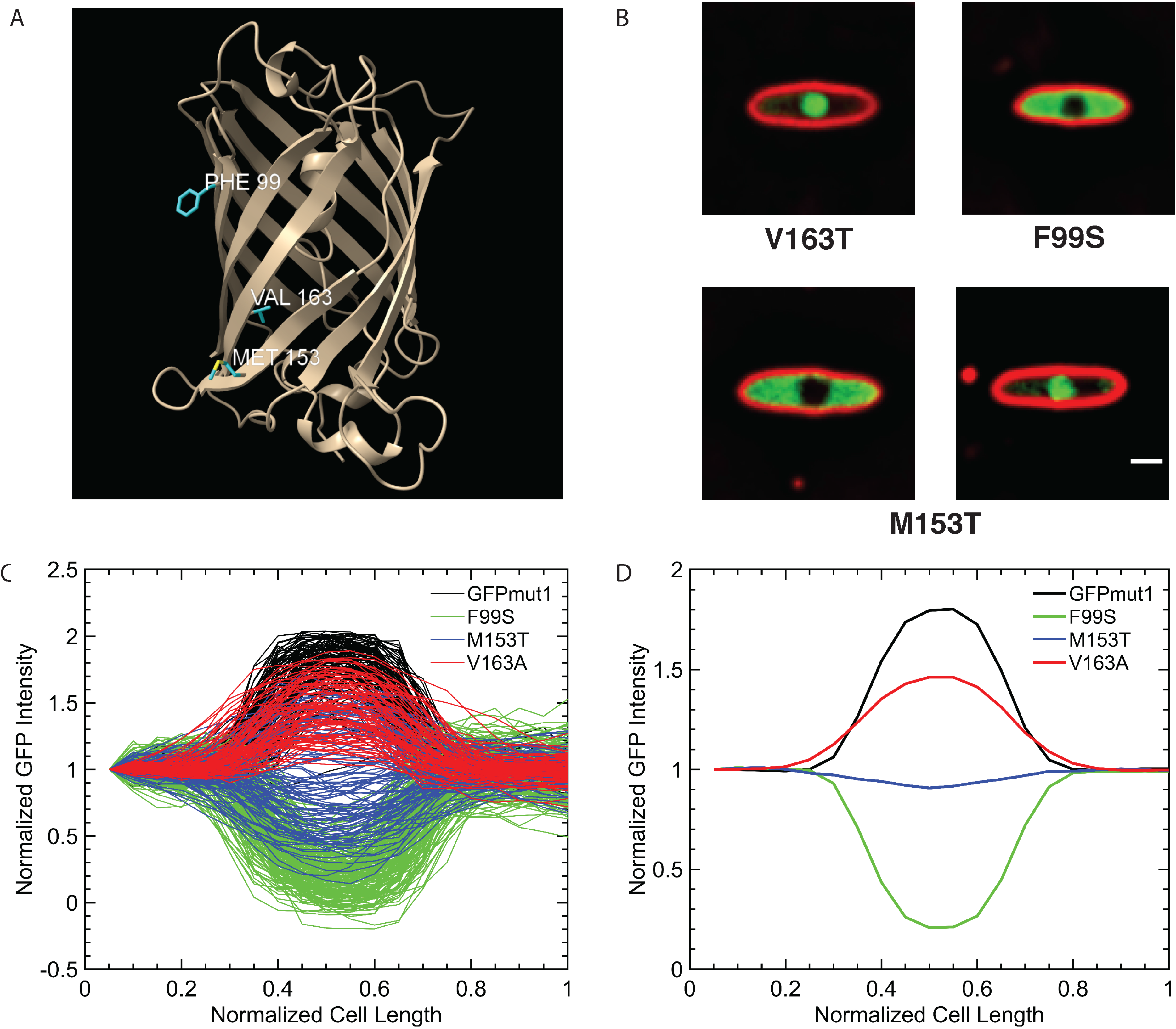
Identification of amino acid residues that alter the nuclear localization of GFPmut1 during PhiKZ infection. A. Three amino acid residues in the GFPmut1 sequence distinguish it from other fluorescent proteins. The mutations are all located on the beta-barrel but two (F99, M153) are on the outer surface while V163 is inside the barrel. B. Three amino acids within GFPmut1 were individually mutated and localized during phage infection. Changing V163 to alanine (V163A) results in nearly 100% localization inside the phage nucleus similar to the unaltered GFPmut1. Changing F99 to serine (F99S) results in nearly 100% cytoplasmic localization during phage infection. M153T appears to localize inside and outside the nucleus in equal measure. Scale bar = 1 micron C. Normalization of GFP intensity in these versions of GFPmut1 was used to quantify the localization of these point mutations in comparison with unaltered GFPmut1. GFPmut1 (n = 111), F99S (n=177), M153T (n=115), V163A (n=82). Each cell expressing GFPmut1 is represented with one black line, showing 100% inclusion into the nucleus. An almost identical phenotype is seen with the red lines representing cells with the V163A mutation. GFPmut1 F99S, shown with green lines, displays GFP intensity outside the nucleus, indicating 100% exclusion. M153T, represented by the blue lines, exhibits both nuclear import and exclusion. D. A plot showing the averages of the individual cells graphed in the left plot. GFPmut1 in black and V163T in red indicate overall inclusion into the nucleus. F99S is represented by green line indicating exclusion. The blue line showing the average of M153T localization profiles is at baseline, showing that on average the protein localizes both inside and outside the nucleus.

We wished to ensure that there were no major structural differences in the ΦKZ nucleus that might explain the differences in permeability. Therefore, we used cryo-EM to visualize the nuclear structures in all three phages. *P. aeruginosa* was infected for 60 minutes with ΦPA3 or ΦKZ, plunge frozen in liquid ethane, and processed for FIB milling and cryo-ET. The process was also performed on *P. chlororaphis* cells infected with 201Φ2-1. We found that the subcellular organization and phage nucleus structure of cells infected with ΦKZ infected cells (Fig 3A, B) were identical to cells infected with 201Φ2-1 (Fig 3C) and ΦPA3 (Fig 3D). The protein shells of all three nuclei formed an unstructured, largely continuous border with a thickness of approximately 5nm (Fig 3). Phage at various stages of maturation were observed, including capsids attached to the side of the nucleus that were either empty or filled with viral DNA, as well as phage tails, some of which were attached to capsids. Bacterial ribosomes were clearly excluded from the phage nucleus as in cells infected with ΦPA3 and 201Φ2-1. These results confirm and extend our previous microscopy experiments (11, 12). Despite the differences in their ability to import GFPmut1, we could discern no obvious differences in the structure of the shell or replication and assembly pathway between these three phages.

**Fig 3:**
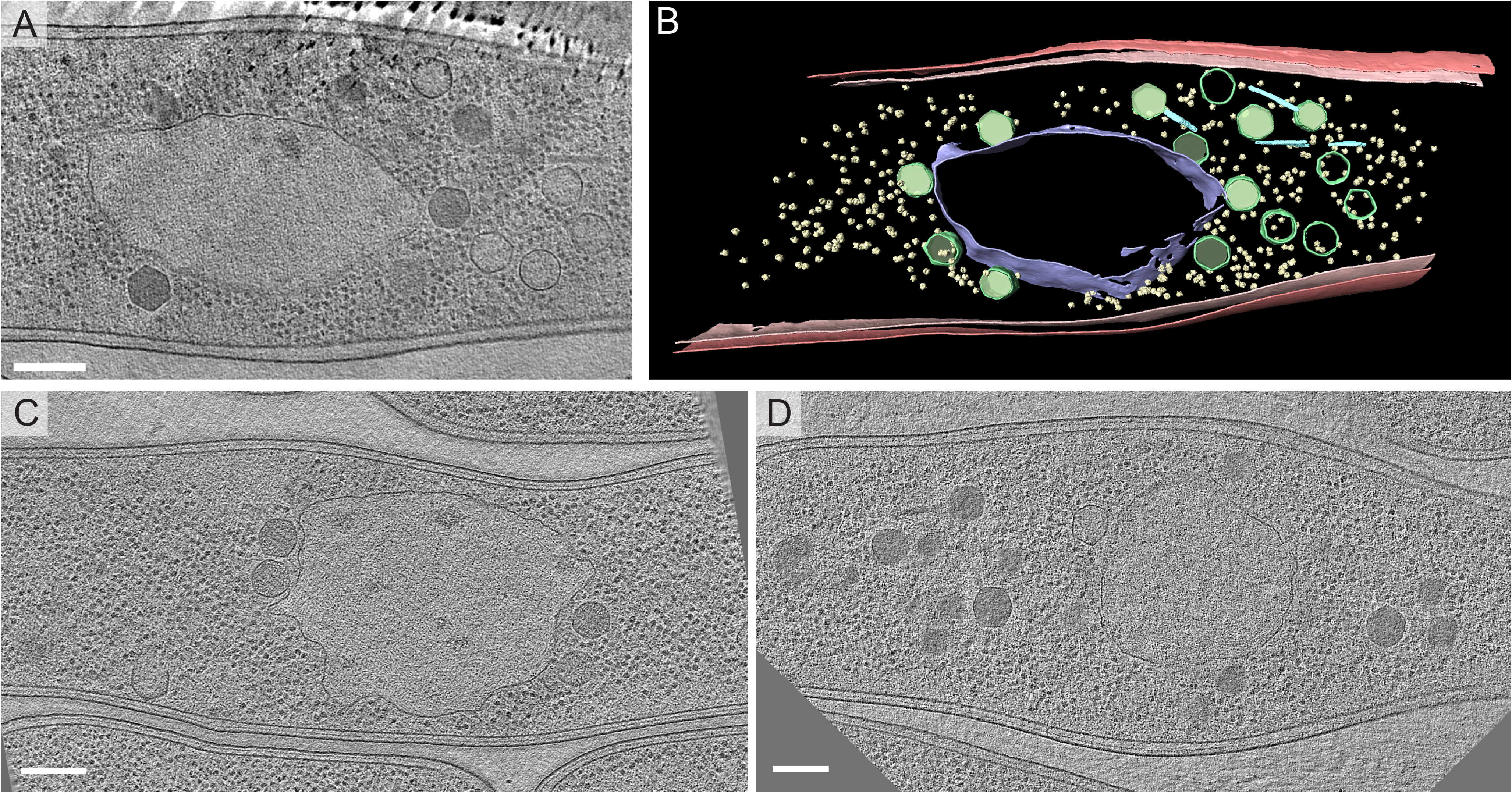
Cryo-EM Tomogram of a *Pseudomonas aeruginosa* cell infected with ΦKZ. A. Slice through a tomogram of cryo-focused ion beam–thinned ΦKZ phage-infected *P. aeruginosa* cell at 60 minutes post infection. Scale bar, 200nm. B. Segmentation of the ΦKZ tomogram shown in (C). The phage nucleus border is shown in darker blue. Bacterial ribosomes are yellow. Phage capsids are green and phage tails are cyan blue. The bacterial cell membrane is shown as red and pink. C. Slice through a tomogram of cryo-focused ion beam–thinned 201Φ2-1 phage-infected *P. chlororaphis* cell at 60 minutes post infection. Scale bar, 200nm. D. Slice through a tomogram of cryo-focused ion beam–thinned 201ΦPA3 phage-infected *P. aeruginosa* cell at 60 minutes post infection. Scale bar, 200nm.

Knowing that GFPmut1 alone was transported into the ΦKZ nucleus, we attempted to test the ability of this fluorescent protein to ferry other proteins into the compartment. As shown previously using cryo-EM and fluorescence microscopy, host bacterial ribosomes are excluded from the nucleus, including the ribosomal subunit L28 tagged with mCherry (12) (Fig 4A). However, tagging the same ribosomal protein with GFPmut1 resulted in its localization inside the nucleus (Fig 4A). Cryo-EM indicated that the ΦKZ tails localized in the cytoplasm (Fig 3A, B). When tagged with sfGFP the major tail protein gp146 formed puncta outside the nucleus; but strikingly, when fused to GFPmut1, gp146 localized inside the nucleus (Fig 4B). GFPmut1 fusion to proteins from other phages were also transported into the ΦKZ nucleus. PA3PhuZ tagged with mCherry formed filaments in the cytoplasm of a cell infected with ΦKZ. However, when fused to GFPmut1, PA3PhuZ localized inside the ΦKZ phage nucleus (Fig 4C).

**Fig 4:**
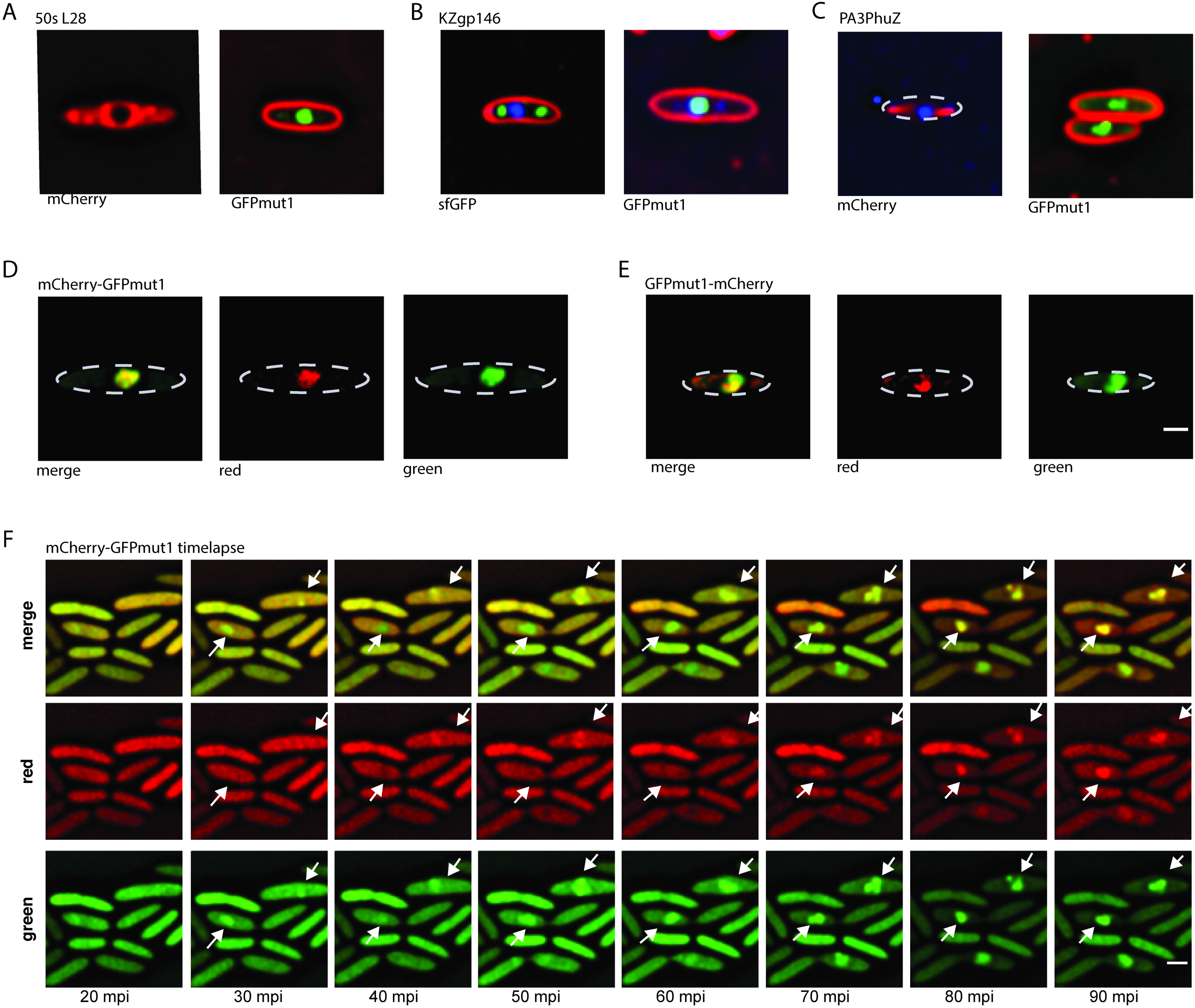
GFPmut1 nuclear localization is dominant in hybrid fusion proteins. Scale bar = 1 micron. A. GFPmut1 fused to host 50s ribosomal subunit L28 localizes inside the phage nucleus while a fusion of the same protein to mCherry localizes in the cytoplasm. B. GFPmut1 fused to ΦKZ tail protein gp146 is mislocalized inside the phage nucleus while a fusion of the same protein to sfGFP shows it localizes as puncta in cytoplasm. C. GFPmut1 fused to ΦPA3 PhuZ protein is seen inside the phage nucleus while a fusion of the same protein to sfGFP forms filaments in the cytoplasm. D. A fusion of mCherry-GFPmut1 shows both proteins are fluorescent inside the phage nucleus. E. A fusion of GFPmut1-mCherry shows both proteins fluoresce inside the phage nucleus after infection. The two fusions in (D) and (E) indicate that GFPmut1 can be fused at both the N and C terminus and retain nuclear targeting. The ability of mCherry to fluoresce indicates that the protein is folded and functional. F. Timelapse of mCh-GFPmut1 shows that both proteins are diffuse in the cytoplasm before infection but move into the nucleus as infection progresses. White arrows indicate two nuclei.

Although the results above suggested that GFPmut1 was able to target proteins to the nucleus, we could not be sure the entire fusion was imported. It remained possible that only GFPmut1 was transported after the target protein was cleaved off. To determine if a tagged protein was imported along with GFPmut1, we created fusions of mCherry at either the N or C terminal ends of GFPmut1 (mCherry-GFPmut1 and GFPmut1-mCherry). This also allowed us to determine if the localization of the target protein was affected by the position of the GFPmut1 fluorescent tag. Both fusions were transported into the nucleus, indicating that fusions to either terminus resulted in nuclear targeting (Fig 4D, 4E). In time-lapse microscopy of cells expressing mCherry-GFPmut1, both green and red fluorescent signals were visible in the cytoplasm before infection. After infection, both green and red fluorescence moved into the nucleus over time, demonstrating that both proteins are transported into the nucleus where they both remain folded (Fig 4F, white arrows). Altogether, our results demonstrate that GFPmut1 can be used to target a fully folded, functional protein to the phage nucleus.

We next attempted to use GFPmut1 to import proteins that might be useful for gene editing. Previous attempts to circumvent the phage nucleus barrier and edit phage genomes relied upon fusing the nuclear targeted RecA-like protein KZgp152 to CRISPR-cas enzymes of interest (26). The RecA-like protein fusion successfully imported a restriction enzyme but failed to import Cas9 (26). Therefore, we tested the ability of GFPmut1 to import three proteins from different CRISPR-cas systems: Cas3, Cas9, and Cas13 (27–31). When fused to sfGFP, all three localized in the cytoplasm, indicating that all three of these proteins are normally excluded from the nucleus (Fig 5A, C, E). In contrast, tagging them with GFPmut1 targeted them into the nucleus (Fig 5B, D, F). These results support our previous hypothesis that the phage nucleus provides a physical barrier that protects phage DNA from endogenous host cell nucleases. In addition, we also examined the host protein SbcB, a single-stranded DNA nuclease that likely functions in host DNA recombination and repair (32, 33). Like the CRISPR-cas proteins, the SbcB-sfGFP fusion localized in the cytoplasm and was excluded from the phage nucleus (Fig 5G). When SbcB was fused to GFPmut1, fluorescence was observed inside the phage nucleus (Fig 5H). Cells expressing sbcB-GFPmut1 also showed misshapen phage nucleoids compared to the smoother shape of the DNA inside infected cells expressing sbcB-sfGFP (Fig 5H). Quantitation of the DAPI intensity of the infection nucleoid in both strains showed that infected SbcB-GFPmut1 cells had a 20% lower average DAPI intensity (approximately 6000 counts, n=187) compared to SbcB-sfGFP expressing cells (average of 7500 counts, n=133), suggesting that internalization of SbcB-GFPmut1 slightly reduces DNA replication or enhances its degradation (Supplemental Fig 2C). When comparing infection of cells expressing SbcB-GFPmut1 to the SbcB-sfGFP counterpart, ΦKZ replication was reduced approximately 10-fold (Supplemental Fig 2A, 2B). Thus, we have shown that GFPmut1 can be used as a nuclear localization tool during ΦKZ infection.

**Fig 5:**
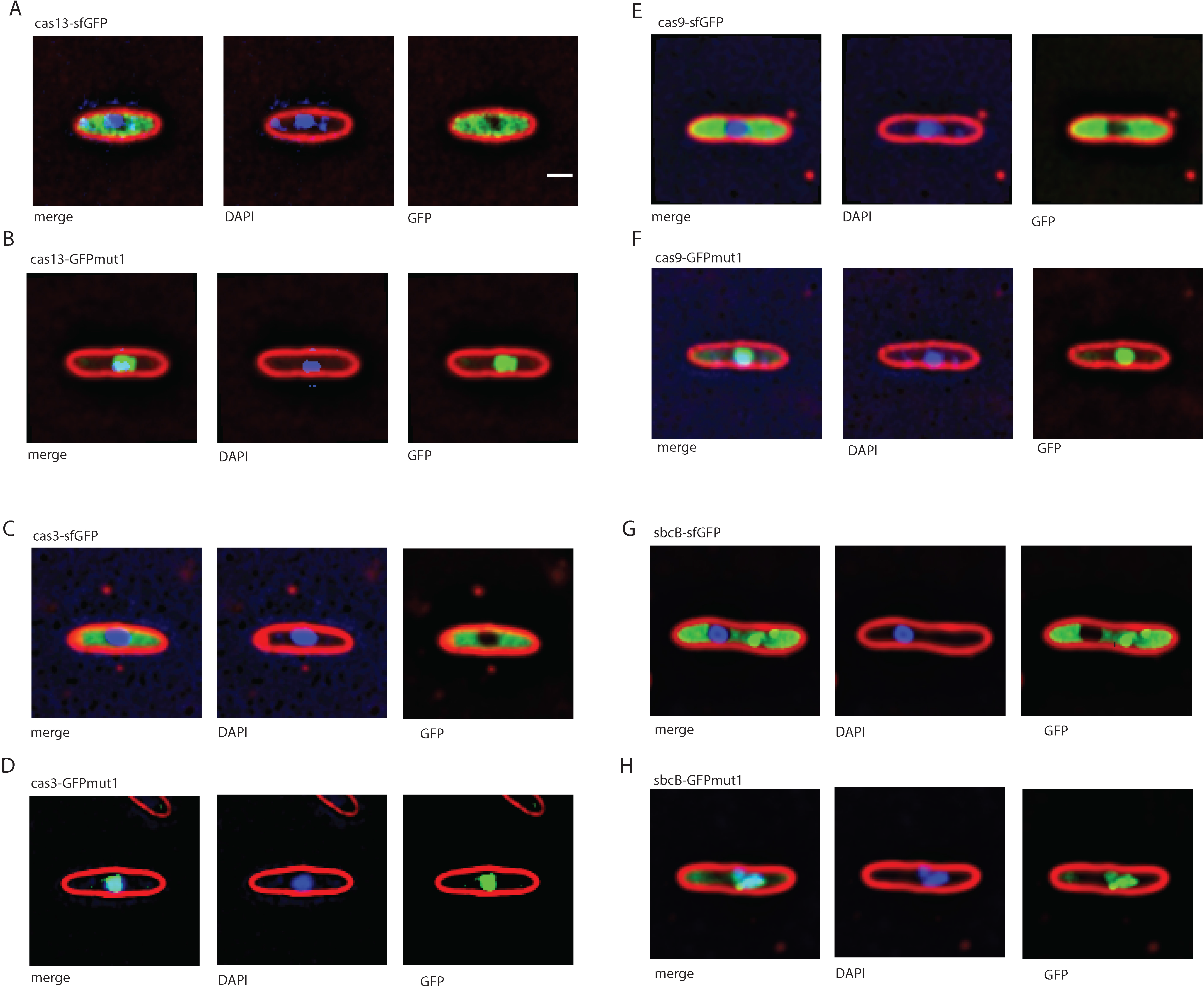
GFPmut1 can be used to artificially import proteins into the ΦKZ nucleus, even those that are detrimental to phage reproduction. Scale bar = 1 micron. A. Cas13 fused to sfGFP localizes outside the phage nucleus. B. Cas13 fused to GFPmut1 localizes inside the phage nucleus. C. Cas3 fused to sfGFP localizes outside the phage nucleus D. Cas3 fused to GFPmut1 localizes inside the phage nucleus. E. Cas12 fused to sfGFP localizes outside the phage nucleus. F. Cas12 fused to GFPmut1 localizes inside the phage nucleus G. SbcB-sfGFP localizes outside the phage nucleus H. SbcB-GFPmut1 localizes inside the phage nucleus with the phage DNA.

## Discussion

Our major finding is that the fluorescent protein GFPmut1, and fusions to it, are transported into the ΦKZ phage nucleus. However, this phenotype is unique to ΦKZ, as GFPmut1 is excluded from the nucleus of the two related phages ΦPA3 and 201Φ2-1. We found these results surprising given the high degree of similarity between these three related jumbo *Pseudomonas* phages (9, 10, 34) and since the cryoEM tomogram of ΦKZ infected cells show a nucleus that is indistinguishable from that of its related phages. The GFPmut1 localization data suggests functional divergence in the selective abilities of these three phage nuclear transport systems.

It remains unclear why GFPmut1 is able to enter the ΦKZ nucleus. Remarkably, a single amino acid change (F99S) completely switches GFPmut1 localization from 100% nuclear to 100% cytoplasmic. One hypothesis is that a protein surface motif is required for recognition by a yet to be identified transport system. The positions of the mutations which have an effect on localization (F99S, M153T) occur on the outer surface of GFPmut1 and imply recognition of the folded structure. The fluorescence of both GFPmut1 and mCherry prior to import supports this idea as well. These results suggest the existence of transport machinery that specifically engages proteins destined for the nucleus. In this model, the surface of GFPmut1 is fortuitously recognized as a substrate and imported by the machinery and the F99S abolishes this interaction.

The unexpected finding of GFPmut1 nuclear targeting raised the possibility that we might be able to use this protein as a convenient way to both label and target proteins to the nucleus of ΦKZ. Understanding which fluorescent proteins are localized outside the ΦKZ nucleus versus which ones are imported is critical for studies of protein localization and will allow us to develop valuable tools for future studies. These results suggest that we can use three different colors of fluorescent proteins (blue, CFP; red, mCherry; or green, sfGFP, GFP^+^, and GFPmut1) to localize proteins during ΦKZ infection, and that we can use GFPmut1 as a tool to specifically target proteins into the nucleus.

Using GFPmut1 to manipulate the ΦKZ nucleus gives us the ability to target and possibly edit phage DNA. Previous attempts to modify the DNA of these large phages have failed (26), most likely because of the physical barrier afforded by the nuclear shell. Our data support our previous hypothesis that a major function of the phage nucleus is protect phage DNA against host defenses, such as CRISPR-cas and restriction enzymes, (12, 26). We now show that GFPmut1 can be used to efficiently circumvent the phage nucleus barrier and target gene editing enzymes into the nucleus, opening up the possibility of genetically manipulating these large phages. Further studies of this targeting phenomenon will also provide insight into the methods utilized by phage ΦKZ for protein sorting. Though the mechanisms used by ΦKZ may differ from the other two phages, determining the specific differences will shed light on the transport systems of the phage nucleus as well as the relationships between these phages. Once we understand the molecular basis of selectivity, we may be able to manipulate it to target proteins to the nuclei in the other phages as well.

## Materials and Methods

### Strain, growth condition, and bacteriophage preparation

*Pseudomonas chlororaphis* strain 200-B was grown on Hard Agar (HA) containing 10 g Bacto-Tryptone, 5 g NaCl, and 10 g agar in 1L ddH_2_O and incubated at 30°C overnight (35). *Pseudomonas aeruginosa* strains PA01 and PA01-K2733 (pump-knockout strain) were grown on Luria-Bertani (LB) media containing 10g Bacto-Tryptone, 5g NaCl, 5g Bacto-yeast extract in 1L ddH_2_O and incubated at 37°C overnight. Lysates for phages 201Φ2-1, ΦPA3, and ΦKZ were made by infecting 5mL of host cultures in early log stage (OD600 = 0.2-0.3) with 500μl of high titer lysate and rolling overnight at 30°C. The phage lysates were then clarified by centrifugation at 15,000 rpm for 10 minutes and syringe filtered through a 0.45 micron filter before storage at 4 °C.

### Plasmid constructions and bacterial transformation

Fluorescent-tagged phage proteins were constructed with the pHERD30T vector as a backbone (36). Phage genes were PCR amplified from phage lysates then ligated into the pHERD30T backbone via isothermal assembly. The assemblies were electroporated into DH5α *E. coli* and plated on LB supplemented with gentamycin sulfate (15μg/mL). Constructs were confirmed with sequencing and subsequently electroporated into either *P. chlororaphis* strain 200-B, *P. aeruginosa* strains PA01, and/or PA01-K2733. *P. chlororaphis* strain was grown on HA supplemented with gentamycin sulfate (25μg/mL) and *P. aeruginosa* strains PA01 and PA01-K2733 were grown on LB supplemented with gentamycin sulfate at 300 μg/mL or 15μg/mL, respectively. See Supplemental Table 1 for a list of plasmids and strains.

### Phage titers

Bacterial cultures were grown in LB Gent 15 liquid broth to late log. 0.5mL culture was then mixed with 4.5mL 0.35% LB top agar and 25uL 20% arabinose (for 0.1% induction) and the mixture poured onto a LB Gent 15 plate. After the top agar lawn had solidified, 5uL of 10x serial dilutions were spotted onto the lawn and the plate was incubated at 30deg overnight.

### Fluorescent Microscopy

The bacterial cells were grown on 1% agarose pads in glass well slides, containing 25% LB, 1ug/mL FM4-64, 1ug/mL DAPI, and 0.1-0.5% arabinose to induce protein expression at desired levels. These pad slides were incubated at 30°C for 3 hours in a humid chamber. For infection beginning at timepoint 0, 5-10 μl of high-titer lysate (10^10^ pfu/ml) was added to pads then incubated again at 30°C. At desired time points after phage infection, a coverslip was put on the slide and fluorescent microscopy performed.

Cells were visualized on an Applied Precision DV Elite optical sectioning microscope with a Photometrics CoolSNAP-HQ2 camera (Applied Precision/GE Healthcare) was used to visualize the cells. For static images, the cells were imaged for at least 8 stacks from the focal plane with 0.15 μm increments in the Z-axis and, for time-lapse imaging, the cells were imaged from a single stack at the focal plane for desired length of time with selected intervals with Ultimate Focusing mode. Microscopic images were deconvolved using SoftWoRx v5.5.1. Image analysis and processing were performed in Fiji.

### Tomography Sample Preparation and Data Acquisition

Infection of *P. chlororaphis* with 201Φ2-1 and *P. aeruginosa* cells with phages ΦKZ and ΦPA3 was done as indicated above. At 70 minutes post infection, cells were scraped off from the surface of the pad using ¼ LB media. 7 μl of cells were deposited on holey carbon coated QUANTIFOIL® R 2/1 copper grids that were glow discharged using Pelco easiGlow^TM^ glow discharge cleaning system. Manual blotting from the side of the grid opposite to the cells using Whatman No. 1 filter paper removed excess liquid such that cells form a monolayer on the surface of the grid. Cells were then plunge-frozen in a mixture of ethane/propane using a custom-built vitrification device (Max Planck Institute for Biochemistry, Munich). Grids were then mounted into modified FEI Autogrids^TM^ to avoid any mechanical damage to the delicate grids during subsequent transfer steps. Then, these clipped grids were transferred into Scios (Thermo Fisher Scientific, formerly FEI), a dual-beam (cryo-FIB/SEM) microscope equipped with a cryogenic stage. Thin sections of 100-250 nm, or lamellae, were prepared as previously described in Chaikeeratisak et al., 2017 containing 10-12 cells each. Tilt-series were collected from typically −65° to +65° with 1.5° or 2° tilt increments using SerialEM^4^ in a 300-keV Tecnai G2 Polara microscope (FEI) equipped with post-column Quantum Energy Filter (Gatan) and a K2 Summit 4k × 4k direct detector camera (Gatan). Images were recorded at a nominal magnification of 34,000 with a pixel size of 0.61 nm. The dose rate was set to 10-12 e/physical pixel at the camera level. Frame exposure was set to 0.1 seconds, with a total exposure in a frame set to be determined by an algorithm targeting an average count number. The total dose in a tomogram was typically ~100-120 e/A^2^ with a defocus of −5 μm. The dataset for this study consists of 16 tomograms from 7 FIB-milled lamellas. Reconstruction of tilt-series was done in IMOD (37) using patch tracking method. Semi-automatic segmentation of the membranes was done using TomoSegMemTV (38) an open-source software based on tensor voting, followed by manual segmentation with Amira software (FEI Visualization Sciences Group).

### Point mutation graph

PA01 cells infected by ΦKZ were imaged 60 to 70 minutes post infection with DAPI staining. Infected cells were identified by the presence of a bright, circular DAPI stain in the center of the bacterial cells corresponding to the presence of phage DNA within the phage nucleus. ImageJ (imagej.nih.gov/ij) was used to bisect infected cells and obtain GFP intensity profiles along their lengths. Each of these intensity profiles were normalized by the length of the cell and normalized again to the GFP intensity at the initial measured end of the cell. Intensity profiles were plotted per cell as well as averaged.

### PDB structure of GFPmut1

The amino acid structure for GFPmut1 was used with the Phyre2 Protein Fold Recognition Server (www.sbg.bio.ic.ac.uk/phyre2) to obtain an estimated structure for GFPmut1 from pSG1729. This sequence differs from EGFP structure 2Y0G by substitutions V1M, L195S and L232H. The resulting structure was viewed with ChimeraX (www.rbvi.ucsf.edu/chimerax). Alignment of fluorescent proteins was made using Clustal Omega (https://www.ebi.ac.uk/Tools/msa/clustalo/)

### DAPI quantification

Images of individual infected cells were cropped using ImageJ. A mask of the phage nucleus was generated using Otsu’s method in Matlab 2017b and the mean DAPI fluorescence was calculated from the raw image intensity within the region of the mask. The complementary image to the mask was used to estimate background fluorescence.

### Growth Curves

Bacterial cultures were grown to late log and then diluted to OD600 0.1. The diluted cultures were induced to 0.1% arabinose and 100uL was aliquoted into each well of 96 well plates. 10uL of phage dilutions were added to appropriate wells. The plate was incubated in the Tecan Infinite M200, shaking, at 30degrees Celsius and OD600 was measured every 10 minutes for 360 cycles (6 hours).

## Acknowledgments

This research was supported by National Institutes of Health grants GM104556 (J.P.) and GM129245 (J.P. and E.V.). We also used the UC San Diego Cryo-EM Facility which is partially supported by a gift from the Agouron Institute and NIH grants to T.S. Baker.

---

**Supplemental Fig 1: A chart showing the amino acid modifications of GFP variants over time**.

**Supplemental Fig 2: sbcb-GFPmut1 shows a small reduction in phage reproduction**.

A. ΦKZ phage titer on a lawn of *Pseudomonas aeruginosa* expressing *sbcB-GFPmut1*. Titer, calculated at 2 × 10^11^ pfu/mL is reduced approximately 10-fold compared to (B).

B. ΦKZ phage titer on a lawn of *Pseudomonas aeruginosa* expressing *sbcB-sfGFP*. Titer is calculated as approximately 2 × 10^12^ pfu/mL.

C. A histogram of DAPI (DNA stain) intensity indicates that cells expressing *sbcB-* mut1 (blue columns, n=187) have lower intensity, compared to cells expressing sbcB-sfGFP (n=133). This suggests that DNA concentration is reduced by the presence of the host nuclease inside the phage nucleus.

## References

1. Hegde RS, Bernstein HD. The surprising complexity of signal sequences. Trends Biochem Sci. 2006;31(10):563–71.

2. Martoglio B, Dobberstein B. Signal sequences: more than just greasy peptides. Trends Cell Biol. 1998;8(10):410–5.

3. Green ER, Mecsas J. Bacterial Secretion Systems – An overview. Microbiol Spectr. 2016;4(1).

4. Chatzi KE, Sardis MF, Economou A, Karamanou S. SecA-mediated targeting and translocation of secretory proteins. Biochim Biophys Acta. 2014;1843(8):1466–74.

5. Patel R, Smith SM, Robinson C. Protein transport by the bacterial Tat pathway. Biochim Biophys Acta. 2014;1843(8):1620–8.

6. Tsirigotaki A, De Geyter J, Sostaric N, Economou A, Karamanou S. Protein export through the bacterial Sec pathway. Nat Rev Microbiol. 2017;15(1):21–36.

7. Thomas JA, Rolando MR, Carroll CA, Shen PS, Belnap DM, Weintraub ST, et al. Characterization of Pseudomonas chlororaphis myovirus 201varphi2-1 via genomic sequencing, mass spectrometry, and electron microscopy. Virology. 2008;376(2):330–8.

8. Serwer P, Hayes SJ, Zaman S, Lieman K, Rolando M, Hardies SC. Improved isolation of undersampled bacteriophages: finding of distant terminase genes. Virology. 2004;329(2):412–24.

9. Monson R, Foulds I, Foweraker J, Welch M, Salmond GP. The Pseudomonas aeruginosa generalized transducing phage phiPA3 is a new member of the phiKZ-like group of ‘jumbo’ phages, and infects model laboratory strains and clinical isolates from cystic fibrosis patients. Microbiology. 2011;157(Pt 3):859–67.

10. Mesyanzhinov VV, Robben J, Grymonprez B, Kostyuchenko VA, Bourkaltseva MV, Sykilinda NN, et al. The genome of bacteriophage phiKZ of Pseudomonas aeruginosa. J Mol Biol. 2002;317(1):1–19.

11. Chaikeeratisak V, Nguyen K, Egan ME, Erb ML, Vavilina A, Pogliano J. The Phage Nucleus and Tubulin Spindle Are Conserved among Large Pseudomonas Phages. Cell Rep. 2017;20(7):1563–71.

12. Chaikeeratisak V, Nguyen K, Khanna K, Brilot AF, Erb ML, Coker JK, et al. Assembly of a nucleus-like structure during viral replication in bacteria. Science. 2017;355(6321):194–7.

13. Erb ML, Kraemer JA, Coker JK, Chaikeeratisak V, Nonejuie P, Agard DA, et al. A bacteriophage tubulin harnesses dynamic instability to center DNA in infected cells. Elife. 2014;3.

14. Zehr EA, Kraemer JA, Erb ML, Coker JK, Montabana EA, Pogliano J, et al. The structure and assembly mechanism of a novel three-stranded tubulin filament that centers phage DNA. Structure. 2014;22(4):539–48.

15. Kraemer JA, Erb ML, Waddling CA, Montabana EA, Zehr EA, Wang H, et al. A phage tubulin assembles dynamic filaments by an atypical mechanism to center viral DNA within the host cell. Cell. 2012;149(7):1488–99.

16. Chaikeeratisak V, Khanna K, Nguyen KT, Sugie J, Egan ME, Erb ML, et al. Viral Capsid Trafficking along Treadmilling Tubulin Filaments in Bacteria. Cell. 2019;177(7):1771–80.e12.

17. Pedelacq JD, Cabantous S, Tran T, Terwilliger TC, Waldo GS. Engineering and characterization of a superfolder green fluorescent protein. Nat Biotechnol. 2006;24(1):79–88.

18. Cormack BP, Valdivia RH, Falkow S. FACS-optimized mutants of the green fluorescent protein (GFP). Gene. 1996;173(1 Spec No):33–8.

19. Crameri A, Whitehorn EA, Tate E, Stemmer WP. Improved green fluorescent protein by molecular evolution using DNA shuffling. Nat Biotechnol. 1996;14(3):315–9.

20. Heim R, Tsien RY. Engineering green fluorescent protein for improved brightness, longer wavelengths and fluorescence resonance energy transfer. Curr Biol. 1996;6(2):178–82.

21. Shaner NC, Campbell RE, Steinbach PA, Giepmans BN, Palmer AE, Tsien RY. Improved monomeric red, orange and yellow fluorescent proteins derived from Discosoma sp. red fluorescent protein. Nat Biotechnol. 22. United States 2004. p. 1567–72.

22. Tsien RY. The Green Fluorescent Protein. Annual Review of Biochemistry. 1998;67:509–44.

23. Ormo M, Cubitt AB, Kallio K, Gross LA, Tsien RY, Remington SJ. Crystal structure of the Aequorea victoria green fluorescent protein. Science. 1996;273(5280):1392–5.

24. Yang F, Moss LG, Phillips GN, Jr. The molecular structure of green fluorescent protein. Nat Biotechnol. 1996;14(10):1246–51.

25. Fukuda H, Arai M, Kuwajima K. Folding of green fluorescent protein and the cycle3 mutant. Biochemistry. 2000;39(39):12025–32.

26. Mendoza SD, Berry JD, Nieweglowska ES, Leon LM, Agard D, Bondy-Denomy J. A nucleus-like compartment shields bacteriophage DNA from CRISPR-Cas and restriction nucleases. bioRxiv. 2018.

27. Ratner HK, Sampson TR, Weiss DS. Overview of CRISPR-Cas9 Biology. Cold Spring Harb Protoc. 2016;2016(12):pdb.top088849.

28. Peters JM, Silvis MR, Zhao D, Hawkins JS, Gross CA, Qi LS. Bacterial CRISPR: accomplishments and prospects. Curr Opin Microbiol. 2015;27:121–6.

29. Charpentier E, Marraffini LA. Harnessing CRISPR-Cas9 immunity for genetic engineering. Curr Opin Microbiol. 2014;19:114–9.

30. Jiang W, Bikard D, Cox D, Zhang F, Marraffini LA. RNA-guided editing of bacterial genomes using CRISPR-Cas systems. Nat Biotechnol. 2013;31(3):233–9.

31. East-Seletsky A, O’Connell MR, Knight SC, Burstein D, Cate JH, Tjian R, et al. Two distinct RNase activities of CRISPR-C2c2 enable guide-RNA processing and RNA detection. Nature. 2016;538(7624):270–3.

32. Phillips GJ, Kushner SR. Determination of the nucleotide sequence for the exonuclease I structural gene (sbcB) of Escherichia coli K12. J Biol Chem. 1987;262(1):455–9.

33. Allgood ND, Silhavy TJ. Escherichia coli xonA (sbcB) mutants enhance illegitimate recombination. Genetics. 1991;127(4):671–80.

34. Cornelissen A, Hardies SC, Shaburova OV, Krylov VN, Mattheus W, Kropinski AM, et al. Complete genome sequence of the giant virus OBP and comparative genome analysis of the diverse PhiKZ-related phages. J Virol. 2012;86(3):1844–52.

35. Thomas JA, Hardies SC, Rolando M, Hayes SJ, Lieman K, Carroll CA, et al. Complete genomic sequence and mass spectrometric analysis of highly diverse, atypical Bacillus thuringiensis phage 0305phi8-36. Virology. 2007;368(2):405–21.

36. Qiu D, Damron FH, Mima T, Schweizer HP, Yu HD. PBAD-based shuttle vectors for functional analysis of toxic and highly regulated genes in Pseudomonas and Burkholderia spp. and other bacteria. Appl Environ Microbiol. 2008;74(23):7422–6.

37. Kremer JR, Mastronarde DN, McIntosh JR. Computer visualization of three-dimensional image data using IMOD. J Struct Biol. 1996;116(1):71–6.

38. Martinez-Sanchez A, Garcia I, Asano S, Lucic V, Fernandez JJ. Robust membrane detection based on tensor voting for electron tomography. J Struct Biol. 2014;186(1):49–61.

