## Supplementary figures and images for "Selective transport of fluorescent proteins into the phage nucleus"

### Supplementary Figure 1

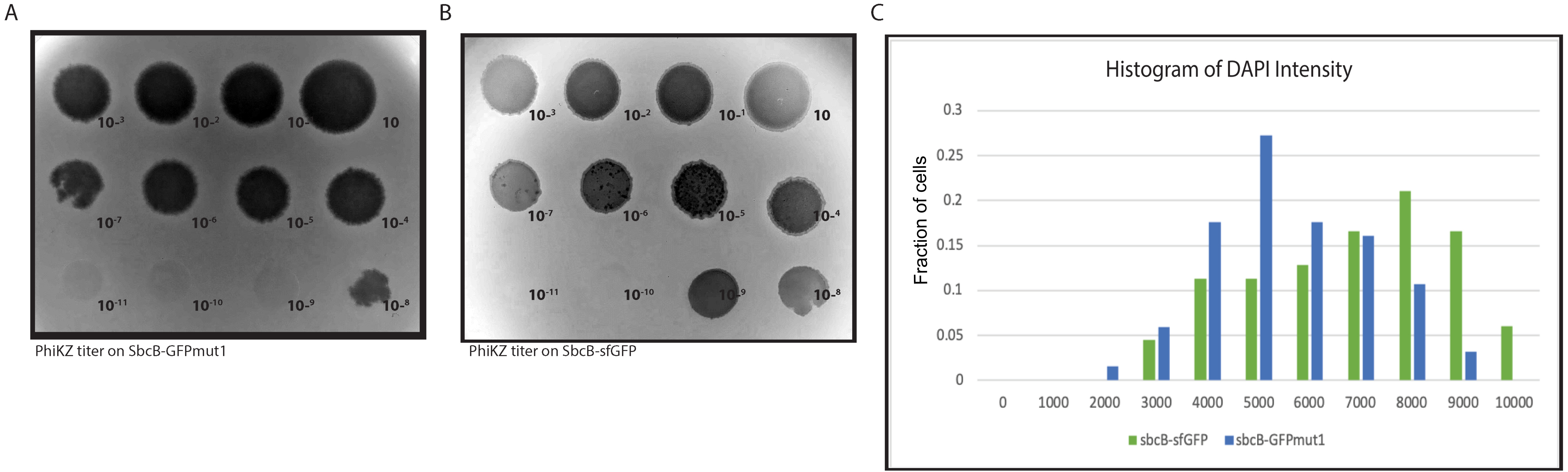

### Supplementary Figure 2

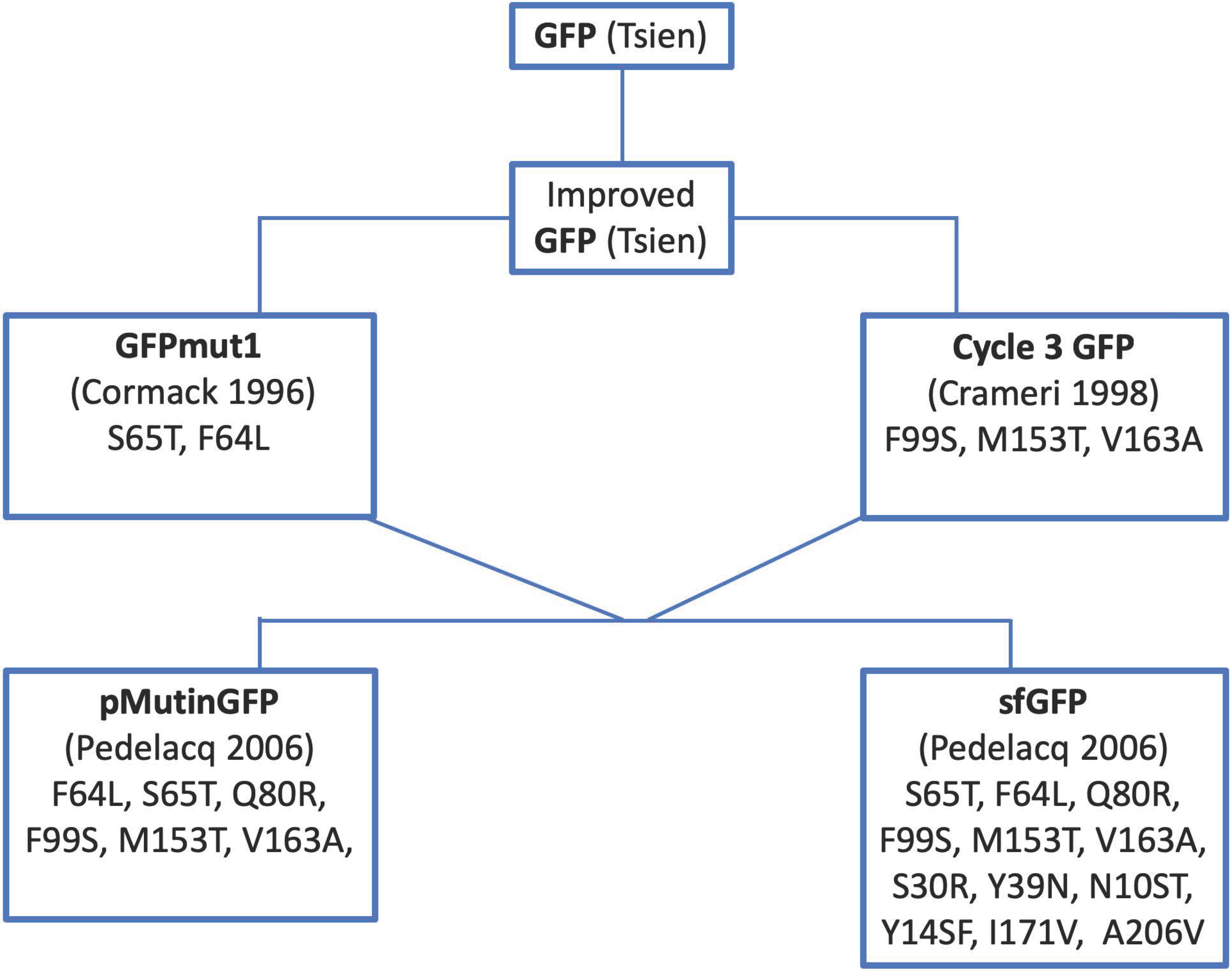
